# Offspring of first-generation hatchery steelhead trout (*Oncorhynchus mykiss*) grow faster in the hatchery than offspring of wild fish, but survive worse in the wild: possible mechanisms for inadvertent domestication and fitness loss in hatchery salmon

**DOI:** 10.1101/2021.09.02.458663

**Authors:** Michael S Blouin, Madeleine C Wrey, Stephanie R. Bollmann, James C. Skaar, Ronald Twibell, Claudio Fuentes

**Author notes:** **Corresponding author** (MSB).

## Abstract

Salmonid fish raised in hatcheries often have lower fitness (number of returning adult offspring) than wild fish when both spawn in the wild. Body size at release from hatcheries is positively correlated with survival at sea. So one explanation for reduced fitness is that hatcheries inadvertently select for trait values that enhance growth rate under the unnatural environment of a hatchery, but that are maladaptive in the wild environment. A simple prediction of this hypothesis is that juveniles of hatchery origin should grow more quickly than fish of wild origin under hatchery conditions, but should have lower survival under wild conditions. We tested that hypothesis using multiple full sibling families of steelhead (*Oncorhynchus mykiss*) that were spawned using either two wild parents (WxW) or two first-generation hatchery (HxH) parents. Offspring from all the families were grown together under hatchery conditions and under semi-natural conditions in artificial streams. HxH families grew significantly faster in the hatchery, but had significantly lower survival in the streams. That we see this tradeoff after only a single generation of selection suggests that the traits involved are under very strong selection. We also considered one possible alteration to the hatchery environment that might reduce the intensity of selection among families in size at release. Here we tested whether reducing the fat content of hatchery feed would reduce the variance among families in body size. Although fish raised under a low-fat diet were slightly smaller, the variation among families in final size was unchanged. Thus, there is no evidence that reducing the fat content of hatchery feed would reduce the opportunity for selection among families on size at release.

## Introduction

Hatchery-reared salmonid fish generally have lower fitness than natural-origin fish when spawning in the wild, and this effect is true even for early-generation hatchery fish (1–7). Here fitness is estimated as relative reproductive success (RRS), the production of returning adult offspring by hatchery fish relative to production by wild fish, when both spawn naturally in the wild. Common garden experiments in steelhead show that the fitness reduction appears to be genetic, rather than simply an environmental effect of the hatchery (1,7,8). That first-generation hatchery steelhead perform worse than natural-origin fish in the wild, but much better as broodstock in the hatchery (9), suggests that the genetic difference results from rapid adaptation to captivity, rather than relaxed natural selection, generalized genomic deterioration, or some other more esoteric explanation (10). Simple adaptation to captivity appears to be a sufficient explanation. Unfortunately, it is not known what traits are under selection in hatcheries, nor the environmental conditions that impose that selection. Understanding those selective pressures might allow hatchery managers to modify hatchery conditions to reduce the rate of domestication, thereby improving the fitness of hatchery fish in the wild.

It is also unclear where in the life cycle the fitness difference between hatchery and wild fish occurs (reviewed in (11)). Studies that estimated RRS by sampling out-migrating smolts and then returning adults found remarkably similar estimates of RRS at the two life stages (2,12). Thus, the fitness loss is already apparent by the time offspring are out-migrating, so we can rule out differential survival at sea as the cause. Therefore, either the hatchery fish themselves have lower breeding ability (fecundity and/or mating competence), or their wild-born offspring have lower survival in the stream environment. Both are plausible explanations, but as yet there is not substantial evidence in favor of either one.

### A simple hypothesis to explain rapid domestication and loss of fitness in hatchery salmon

The size of hatchery salmon at release is positively correlated with their probability of survival at sea (13–18). Therefore, a plausible hypothesis to explain rapid domestication is that hatcheries select for values of physiological or behavioral traits that allow some fish to grow quickly in the unnatural conditions in a hatchery (10,14,19). Such selection might be especially intense on steelhead, which are typically forced to smolting in one year in hatcheries versus the typical two years that they take in the wild (20,21).

The hatchery environment features no predators, an abundance of calorie-dense food, and a very crowded social environment. The values of physiological and behavioral traits that might allow a fish to thrive and grow quickly in that environment would probably be maladaptive in the low-food, predator filled environment of a natural stream. So under this scenario, hatchery fish having those extreme trait values would grow quickly, be large at release, and so survive to return from sea. However, when they pass those genes on to their offspring in the next generation, their offspring would have poor survival in the wild. Thus, under this scenario the fitness difference between hatchery and wild fish would result mainly from differential survival of their offspring, not from differential fecundity or mating ability of the hatchery adults. If the above hypothesis is true, then the offspring of hatchery fish should (1) grow faster under hatchery conditions than the offspring of wild fish, but (2) have slower growth and/or poorer survival under natural conditions. Furthermore, among families within each type of fish there should still be genetic variation for whatever traits are under selection. So a corollary prediction is that, within each fish type, families that grow more quickly in the hatchery should perform worse under natural conditions.

Here we tested the above hypotheses by comparing multiple full-sibling families of steelhead (*Oncorhynchus mykiss)* created using either two wild parents (WxW) or two first-generation hatchery parents (i.e. result of one generation of selection in the hatchery; HxH). We mixed individuals from all the families together and raised them under hatchery conditions and under semi-natural conditions in artificial streams in order to compare their growth and survival in both environments.

### Test of a possible method for reducing adaptation to the hatchery

In addition to testing whether HxH and WxW fish differ in survival or growth in the hatchery vs. wild, we simultaneously tested whether altering the fat composition of feed could alter the performance difference between the two types of fish in the hatchery. The idea here is that pelleted fish food has a much higher fat content than the natural diet. So any physiological adaptation that allows some fish to more efficiently turn that fat into increased body size and condition would be selected for. Thus, reducing the fat content of feed might reduce, or perhaps even reverse, any performance difference seen between high- and low-growth families under normal feed conditions.

Commercial feed developed for hatcheries is unnatural in that it is typically high in fat and protein, calorically rich, and high in marine derived nutrients. This nutrient composition is novel for juvenile steel head, which typically hunt in riffles of streams. Gut content analysis of wild salmon indicates that a large portion of the normal diet is composed of aquatic insect larva (22), which although high in protein, are not high in fat (23). Variation in ability to metabolize high-fat hatchery feed, and in turn, the ability to rapidly convert that feed into somatic growth at the juvenile stage should directly contribute to size and body condition at release. Fat stores in anadromous fish are critical for successful outmigration post-release (24). Thus, in a hatchery rearing environment, fish that have the ability to take advantage of such an abundance of high-fat food would be at a growth advantage, and more likely to be big enough and have the fat reserves to survive when released after only a year of fresh-water rearing. Conversely, a fish well adapted to utilize a high fat diet for rapid growth might be at a disadvantage in the wild where food resources are lower in fat content and far less abundant.

Here we tested whether lowering the fat content of the diet could reduce the variation among families in performance (growth), and/or affect any performance difference observed between HxH and WxW fish. If so, then altering the diet might be one way to reduce the opportunity for selection on whatever traits control growth rate in the hatchery.

## Methods

### Fish stock and spawning

Twenty HxH and 20 WxW full-sibling groups were created using one-to-one pairings of adult steelhead from the Wilson River in Oregon. Returning wild adults and first-generation hatchery adults (created using wild brood stock) were caught in traps and held until spawning during February-March, 2017 at the Oregon Department of Fish and Wildlife’s Trask hatchery (returning hatchery fish are identifiable by their missing adipose fins). Spawning of brood stock occurred once weekly between Feb 22, 2017 and March 14, 2017 with approximately equal numbers of wild- and hatchery-origin pairs spawned on each day. Groups of full sib eggs were reared separately. Eyed eggs were transferred to the OHRC and incubated in Heath trays until yolk absorption. Embryos from the early spawning dates were reared on chilled water to delay development, ensuring that exogenous feeding would begin on the same day for all offspring. Each family group of embryos was reared separately until ponding to ensure that the number of offspring from each family was equal at the start of each experiment.

One family of WxW fish (family C) had very poor survival to hatching, so there were not enough fry to include family C in the hatchery tank growth rate experiment. So we excluded that family and one of the HxH families (family I) in order to have a balanced design of 19 families of each type in the hatchery tanks. We included family I in the stream experiment. We also added a small number of surviving fry from family C in the streams just to see if their performance would be similar to that of the other families, but did not include them in the analysis of survival.

### Experimental set up - streams

The OHRC has four artificial streams that replicate natural channels with pools, riffles, and woody debris in a gravel bed. Water is pumped to each stream from Fall Creek and allowed to flow the length of the stream by gravity. Fish are retained by screens at the end of the stream. The streams are approximately 55 m in length and 7 m in width, and range in depth from around 15 cm in the riffles to 60 cm in the pools. Fish feed on the natural invertebrate fauna that develops in each stream. The streams were loosely covered with shade cloth and bird netting, but some bird predators (kingfishers, *Megaceryle alcyon*) regularly entered the enclosure. There were no large aquatic predators.

On May 15, 2017, unfed fry from each family were released into two of the artificial streams at the OHRC. Sixty individuals from each of the 39 families in the analysis were added to each stream (120 total per family). For family C we only had enough surviving fry to add 30 per stream. On October 28-29, 2017, ODFW staff electroshocked the streams to collect all the surviving fry, which were euthanized in MS-222. Each fry was weighed, measured (fork-length) and sampled for DNA.

Unfortunately, the meta-data associated with each fin sample from the fish in the streams were lost. So all we know is the family identity of each sample, but not which of the two streams it came out of or its body size. So we are able to measure the overall survival of each family as the number of survivors out of 120 siblings, but not the average body size of those survivors. Nevertheless, we can still test two *a priori* hypothesis: (1) WxW families should have higher survival than HxH families in the streams, and (2) within fish type, faster growing families in the hatchery should have poorer survival in the streams.

### Experimental set up – hatchery tanks

On May 15, 2017, unfed fry were transported to Oregon State University’s Aquatic Animal Health Lab, where they were ponded into 400 L, covered outdoor tanks. Thirty fish from each of 38 families (19 HxH and 19 WxW) were added to each 400 L tank (N = 1140 per tank). Fish were fed from May 22^nd^ to November 6^th^, 2017. Food was dispersed by belt auto-feeders, set to deliver food over an 8-hour period daily from first feeding until August 10^th^, 2017. From August 10^th^ to the end of the experiment on November 6^th^, 2017, fish were fed by hand daily because wet weather conditions interfered with auto-feeder function. Fish in each tank were measured every other week in order to monitor growth and adjust feed amounts as needed. Fish were fed according to manufacturer recommendations based on water temperature and average body size. An excess of food was available daily, so fish should have been able to feed to satiation.

We started with 1140 fish per tank in order to mimic standard hatchery densities when fish are first learning to feed. We did not have the option of moving the fish to raceways or larger tanks as they increased in size, so on July 24-26 we randomly euthanized 570 fish from each tank to maintain appropriate stocking densities. On August 10^th^, 2017, we again adjusted stocking densities by subdividing each original tank into two 400 L tanks. Fish were reared in these tanks until they were lethally sampled on November 6, 2017.

### Feed composition experiment

The families were raised in the hatchery using two types of feed in order to test whether reducing the fat content of feed might affect any performance difference between HxH and WxW families in the hatchery, or the variation in growth rate among families within each fish type. If so, then altering feed might be one way to reduce the intensity of selection in the hatchery.

The two dietary lipid treatments were obtained by adding different amounts of marine fish oil to a series of commercial salmonid diets (Bio Vita Starter sizes #0 crumbles, #1 crumbles, #2 crumbles and Bio Clark’s Fry size 1.2 mm) that had not been top-coated with lipid by the manufacturer (Bio Oregon, Inc., Longview, WA). Lipid concentrations in the low lipid diets were approximately 12%. Lipid concentrations in the standard lipid diets were 18-20%, which is similar to those in commercially available salmonid diets.

We started with six 400 L tanks, three replicates of each feed-type treatment. We ended with 12 tanks in total after splitting each tank into two tanks on August 10^th^ in order to maintain appropriate stocking densities. In the data analysis below, we refer to the original 3 tanks per treatment as “tanks” and the two tanks into which we split each tank as “sub-tanks”.

### Genotyping and parentage assignment

At the end of each experiment all fish were matched to their parents by genotyping them at six microsatellite loci (SPAN-B primers; Stephenson et al., 2009)(25). Parentage was assigned using pure exclusion via the SOLOMON program in R (26).

### Statistical analysis: Effects of fish type (HxH vs. WxW) and food treatment (high vs. low fat) on body size in the hatchery

We tested the effect of fish type and food treatment on growth in the hatchery in two ways. Firstly, we used a linear mixed model analysis in SYSTAT 13.2 (Systat software, Inc.) to test and estimate the effects of fish type and food treatment on body size while accounting for the effects of family, tank and sub-tank. Model parameters included the following: intercept, fish type, food treatment, and fish type-by-treatment interaction as fixed effects, and family (nested within fish type), tank (nested within treatment) and sub-tank (nested within tank) as random effects.

Secondly, we also took a simpler approach to asking the same question, which gave similar results, but is easier to interpret graphically. Here we simply pooled all six sub-tanks within each treatment and estimated the average body size of each family within each treatment. Thus the unit of replication here is the family, and we had 19 data points for each fish type within each treatment. We then did a two-way General Linear Model (GLM) in Systat 13.2 using fish-type, treatment and their interaction as parameters. Note that pooling all six sub-tanks may add tank effects to the error term, resulting in a less powerful test for the effects of interest (fish type and treatment), but should not introduce any bias into the results.

### Statistical analysis: Effects of fish type (HxH vs. WxW) on survival in the streams

We first used a Mann-Whitney U-Test to test whether WxW and HxH families differed in number of survivors. We contrasted those results with those obtained from a parametric ANOVA using either raw counts or arcsin square root transformed proportions (arcsin of the square root of the proportion) (27).

### Statistical analysis: Correlation between growth rate of each family in the hatchery and survival of each family in the streams

Our *a priori* hypothesis was that, among families of each type, there would be a negative correlation between survival in the streams and growth rate in the hatchery. We have a single measure of survival for each family, but measured growth in the hatchery under two different conditions – under a high fat diet and under a low fat diet. So for each measure of growth we did an analysis of covariance on survival (counts) using growth rate as the continuous variable and fish type as categorical. We compared additive and non-additive (with interaction term) models via Extra Sum of Squares tests. The additive model was more appropriate in both analyses, and neither interaction was significant, so we report coefficient estimates from the additive model. In addition to the standard linear regression approach, we also performed Poisson and Negative-Binomial regression to directly model the counts while accounting for possible over-dispersion in the data. These analyses were performed in R (Core Team 2021).

### Animal Research

This work was carried out under protocols approved by Oregon State University’s Animal Care and Use Committee (protocol #4704).

## Results

### Effects of fish type and food treatment on growth in the hatchery

The HxH families were, on average, larger than the WxW families under both food treatment conditions (diets), and fish raised under a high-fat diet were larger than those raised under low-fat diet (Fig 1). The mean, standard deviation and coefficient of variation of the average family body sizes from Fig 1 are shown in Table 1. The GLM on family mean sizes showed significant effects of both fish type (P = 0.001) and food treatment (P = 0.015), but no interaction between the two factors (P = 0.709; Table 2). The estimated overall effect sizes were 2.2 mm larger for HxH families and 1.7 mm larger for fish raised under high-fat food.

**Table 1.** Means, standard deviation (SD) and coefficient of variation (CV) of the family means shown in Fig 1, by fish type and diet.

| Combination | Mean | SD | CV |
| --- | --- | --- | --- |
| HxH-High Fat | 127.6 | 5.60 | 0.044 |
| WxW-High Fat | 123.6 | 6.37 | 0.052 |
| HxH-Low Fat | 124.74 | 5.05 | 0.040 |
| WxW-Low Fat | 119.74 | 6.18 | 0.052 |

**Table 2.**
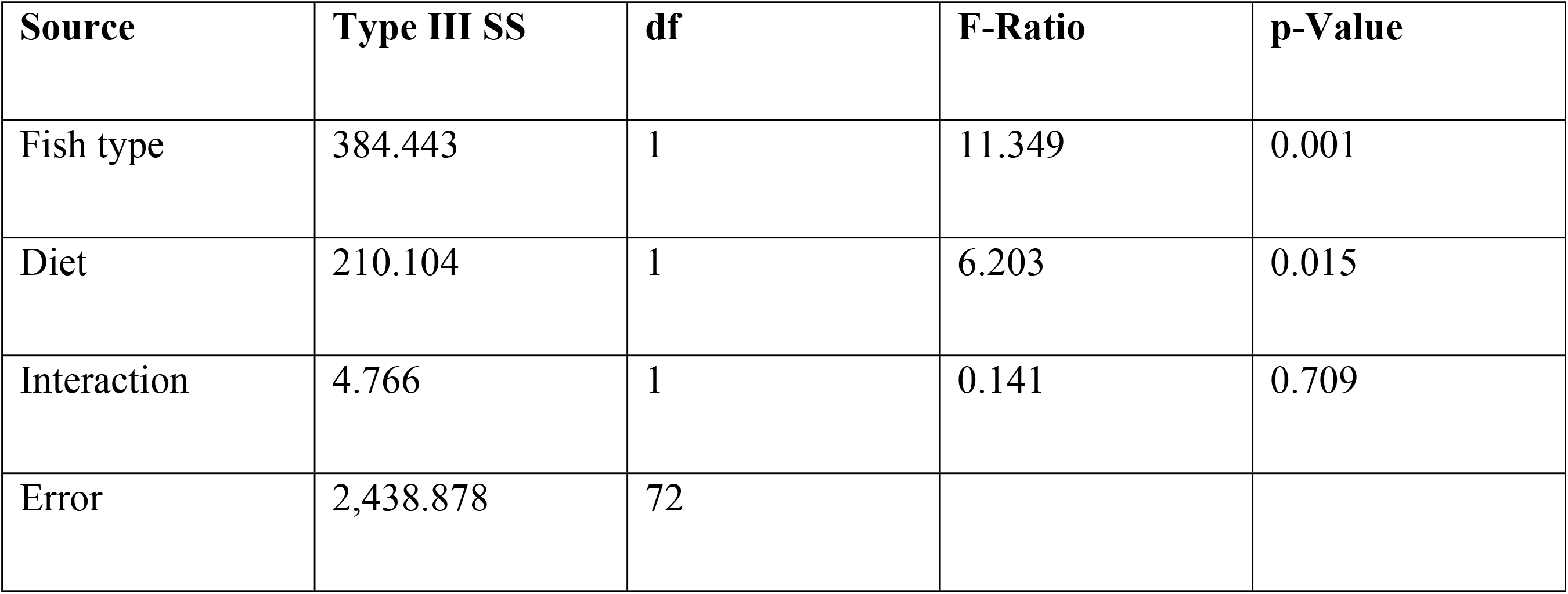
GLM on average family sizes.

**Fig 1.**
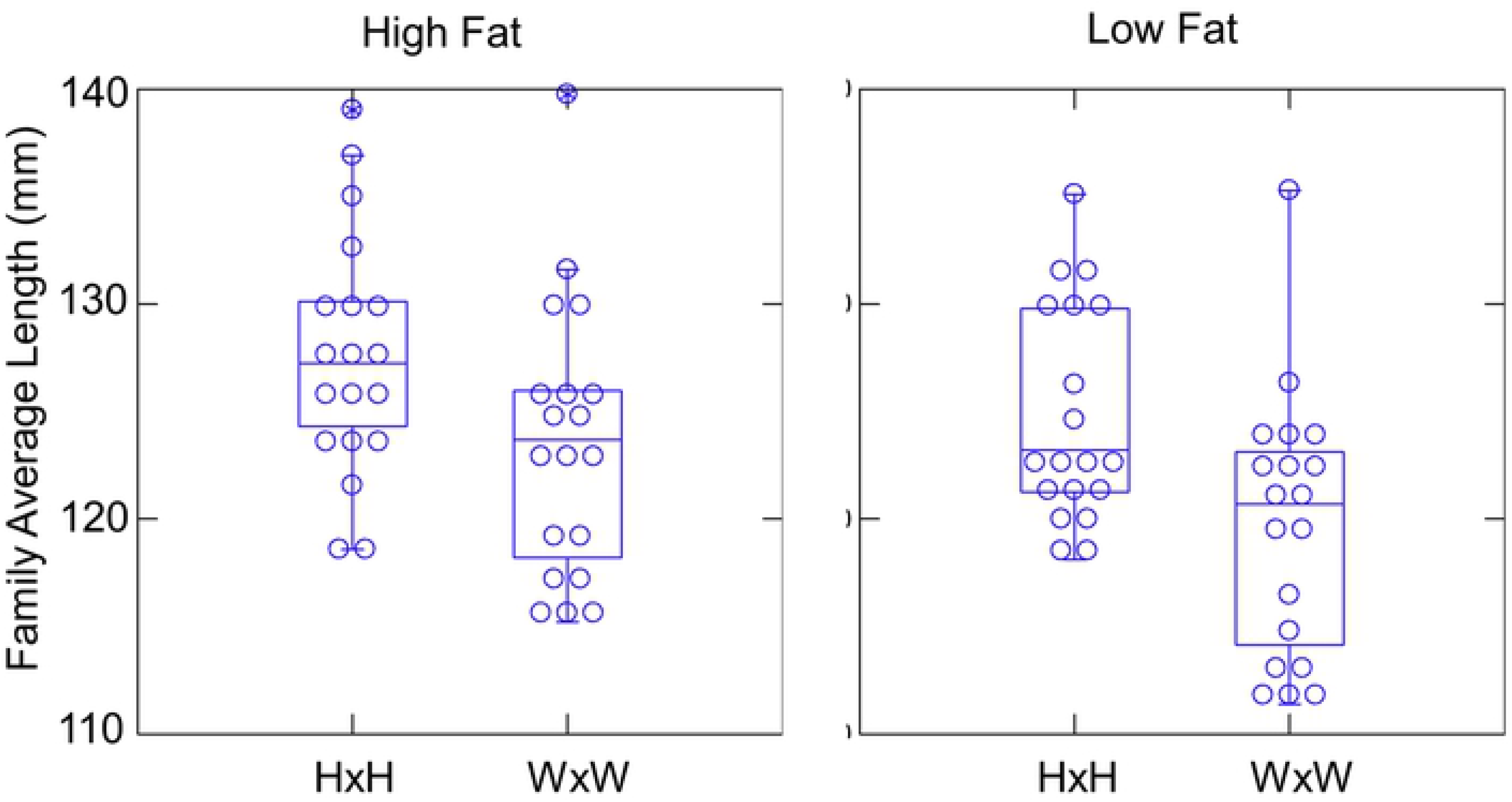
Average body size of each family raised under high fat (High) and low fat (Low) diet, by fish type. HxH families grew faster than WxW families under both treatments. The same data are shown as both dot density plots and as box plots.

The standard deviation and coefficient of variation of length among families changed little between food treatments (Table 1). So there is no evidence that altering the diet reduced the variation in performance among families of either type. Thus, reducing the fat content of food seems unlikely to reduce the opportunity for selection on body size among families.

The linear mixed model analysis on the entire dataset of individual fish also showed that HxH fish grew significantly faster than WxW fish under both food treatments (P = 0.019; Table 3). HxH fish averaged an estimated 5.1 mm larger than WxW fish. Fish raised on the high fat diet were estimated to be, on average, 4.1 mm larger, but this difference was not significant in the mixed model (P = 0.126). There was again no significant fish-type by treatment interaction (P = 0.089), so we conclude that HxH fish outperformed WxW fish similarly under both food conditions. Among the random effects, family explained 25.2% of the total variance in length, while tank and sub-tank explained 0.8% and 6.1% respectively. So, family identity has a very large influence on growth of steelhead in the hatchery, as seen in other studies in which fish were raised under similar conditions (19).

**Table 3.** Type III Tests for Fixed Effects from Linear Mixed Model.

| Effect | Numerator df | Denominator df | F-Ratio | p-Value |
| --- | --- | --- | --- | --- |
| Fish type | 1 | 36 | 6.057 | 0.019 |
| Diet | 1 | 4 | 3.733 | 0.126 |
| Interaction | 1 | 2,822 | 2.887 | 0.089 |

### Survival in the streams: HxH vs WxW families

The number of survivors per family for the 39 families used in the streams analysis ranged from 5 to 36 (out of 120 per family stocked per family initially). We found only a single surviving individual from family C, confirming that family’s *a priori* status as an outlier. WxW families had higher survival than HxH families (Fig 2; Mann-Whitney *U* = 110.5, N_HxH_ = 20, N_WxW_ = 19, *P* = 0.025). A parametric ANOVA using either raw counts or arcsin square root transformed proportions also showed that WxW families had significantly higher survival (P = 0.019 and 0.015 respectively).

**Fig 2.**
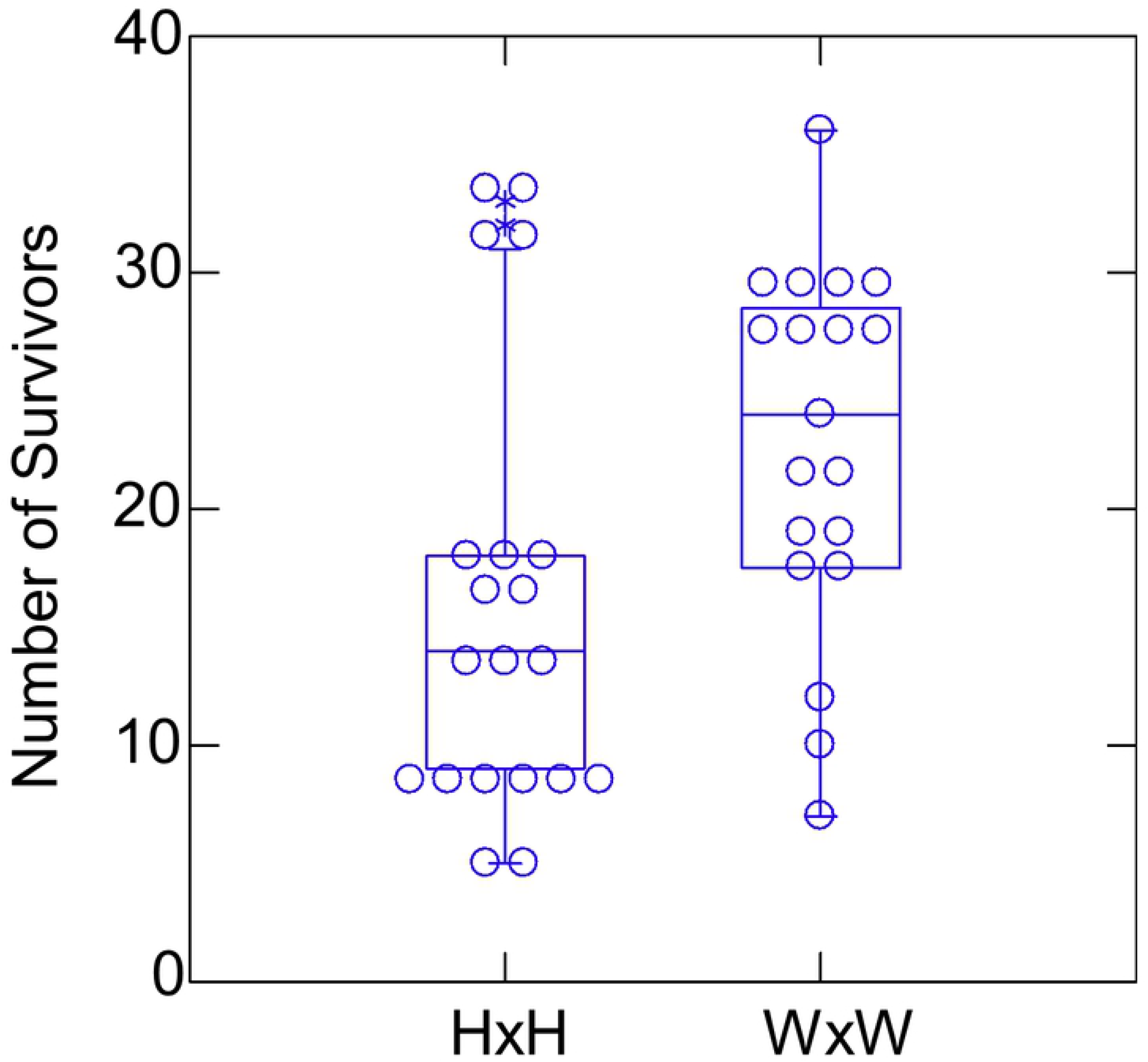
Number of survivors in the streams per family (out of 120 starting individuals per family), broken down by fish type. The same data are shown as both a dot density plot and as a box plot.

### Family correlation between growth rate in the hatchery and survival in the streams

Our prediction was that, within fish type, there would be a negative correlation among families between growth in the hatchery and survival in the streams. Although the overall estimated correlation was negative as expected, it was not significantly different from zero (Fig 3 and Table 4). Poisson and Negative Binomial regression corroborated these results (S1 Table).

**Table 4.** Effect Size Estimates from ANCOVA of Survival vs. Family Average Size

| Coefficient | Estimate | Std Error | p-Value |
| --- | --- | --- | --- |
| (A) Size measured under high fat diet |  |  |  |
| Intercept | 56.39 | 30.38 | 0.072 |
| Size | -0.314 | 0.238 | 0.195 |
| Fish type (WxW) | 5.11 | 2.93 | 0.090 |
| (B) Size measured under low fat diet |  |  |  |
| Intercept | 66.09 | 31.22 | 0.042 |
| Size | -0.399 | 0.250 | 0.119 |
| Fish type (WxW) | 4.37 | 3.01 | 0.156 |

**Fig 3.**
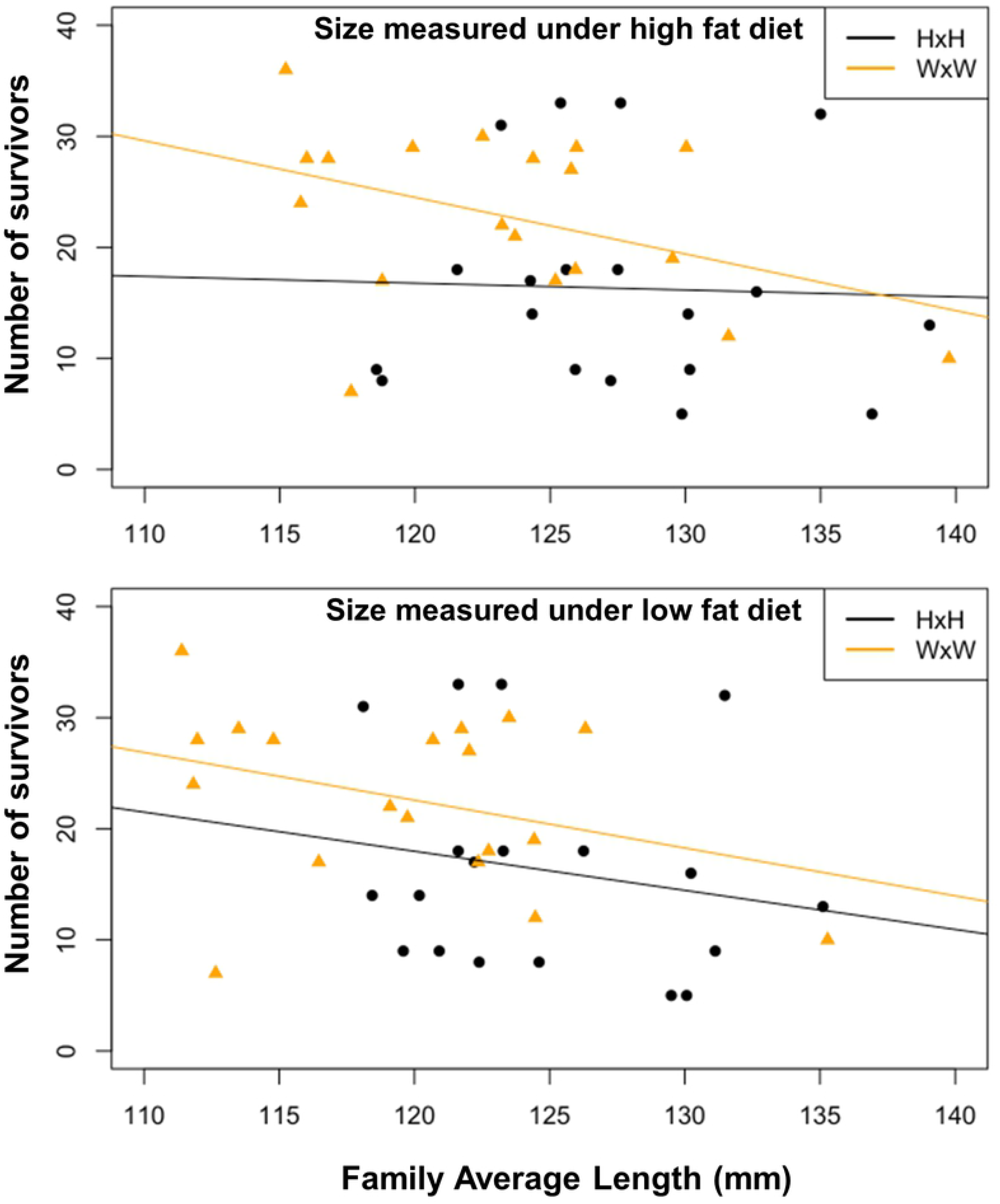
Relationship among families between survival in the streams (counts of survivors) and average family body size measured in the hatchery under a high fat diet (top) or a low fat diet (bottom). In both cases there was no significant interaction between size and fish-type, and the correlation was not significantly different from zero.

## Discussion

In this study the offspring of first-generation hatchery steelhead grew faster in the hatchery, but had lower survival under wild conditions, than the offspring of wild steelhead. These results are consistent with data on lifetime reproductive success (production of returning adult offspring) from another population of steelhead that show clear evidence of adaptation to the hatchery, with tradeoffs in fitness in the wild, after a single generation of selection (data from Hood River steelhead; (9)). That we again see differences after only one generation of selection in the hatchery illustrates how strong the selection pressures must be on whatever traits are involved.

The hypothesis that there exists genetic variation among families for traits that promote fast growth in the hatchery, but with a cost in offspring survival in the wild, predicts that one would also see a negative correlation among families between growth in the hatchery and survival in the wild. We saw no significant relationship, although the trend was in the predicted direction. This hypothesis would be worth testing again to determine if a negative correlation really exists.

These results also support the hypothesis that the reduced fitness of hatchery adults in the wild results from lowered survival of their offspring, rather than only from reduced fecundity or mating ability of the returning hatchery adults themselves. The fact that offspring survival in the wild is probably involved is consistent with the hypothesis that fitness loss involves selection on heritable behavioral or physiological traits of the juveniles in the hatchery. Thus, it remains plausible that one could modify the hatchery to reduce the selection pressures if one could identify the traits under selection and what aspects of hatchery culture cause that strong selection.

One possible way to modify the hatchery would be to change the feed composition to match more closely the nutritional composition of what fish eat in the wild. This suggested modification follows from the hypothesis that hatcheries might select for physiological traits that allow some fish to take advantage of the novel food source (abundant, high-fat). However, our results show that reducing the fat content of feed to more natural levels did not reduce the variation among families, and if anything, exacerbated the performance advantage of the HxH families over the WxW families (Fig 1). So perhaps other traits, such a behavior, are more important, or perhaps other aspects of feed composition are more important. Regardless, there is no evidence that this particular modification to standard hatchery practice would reduce the opportunity for selection on final body size.

## Conclusions

In summary, fish whose parents underwent a single generation of selection in the wild grew faster in the hatchery and had poorer survival in a semi-natural stream. There was a non-significant relationship between family growth in the hatchery and survival in the stream, although the nominal correlation was negative. Reducing the fat content of the fish’s diet had little effect on the variation among families in body size. So altering the fat content of feed does not seem like a promising way to reduce the opportunity for selection among families on size at release in a hatchery. Nevertheless, it remains plausible that there are other aspects of hatchery practice that one could modify to reduce the selection pressure, thereby producing hatchery fish having fitness in the wild that is more similar to that of wild fish.

## Acknowledgments

Thanks to the Oregon Department of Fisheries and Wildlife (ODFW), to Jen Krajcik, Joseph O’Neill and staff at the Oregon Hatchery Research Center, to Ruth Milston Clements and staff at OSU’s Aquatic Animal Heath Lab, to staff at the Trask hatchery, and to Franco Felix for invaluable assistance in obtaining eggs and raising fish for these experiments. Thanks to Stan Cates and to OSU’s Center for Quantitative Life Sciences for help with genotyping. The findings and conclusions in this article are those of the authors and do not necessarily represent the views of the U.S. Fish and Wildlife Service.

## Supporting Information

**S1 Table. Coefficients from Negative Binomial and Poisson regressions of average size of each family against survival (counts) of each family.**

